# The mitochondrial genome of the red icefish (*Channichthys rugosus*) casts doubt on its species status

**DOI:** 10.1101/2022.06.27.497707

**Authors:** Moritz Muschick, Ekaterina Nikolaeva, Lukas Rüber, Michael Matschiner

## Abstract

Antarctic notothenioid fishes are recognized as one of the rare examples of adaptive radiation in the marine system. Withstanding the freezing temperatures of Antarctic waters, these fishes have diversified into over 100 species within no more than 10-20 million years. However, the exact species richness of the radiation remains contested. In the genus *Channichthys*, between one and nine species are recognized by different authors. To resolve the number of *Channichthys* species, genetic information would be highly valuable; however, so far, only sequences of a single species, *C. rhinoceratus*, are available. Here, we present the nearly complete sequence of the mitochondrial genome of *C. rugosus*, obtained from a formalin-fixed museum specimen sampled in 1974. This sequence differs from the mitochondrial genome of *C. rhinoceratus* in no more than 27 positions, suggesting that the two species may be synonymous.

## 1 Introduction

The diversification of fishes of the perciform suborder Notothenioidei in Antarctic waters is a rare example of adaptive radiation in the marine environment (Clarke & Johnston, 1996; Eastman, 2005; Matschiner et al., 2015; Near et al., 2012; Rüber & Zardoya, 2005). The radiating group of notothenioid fishes is composed of five families (Nototheniidae, Harpagiferidae, Artedidraconidae, Bathydraconidae, and Channichthyidae; jointly called ”Cryonotothenioidea” (Near et al., 2015)) that together include over 100 species, distributed primarily on the shelf areas surrounding Antarctica and sub-Antarctic islands (Eastman & Eakin, 2021; Gon & Heemstra, 1990). The validity of most species within this radiation is well established and in many cases corroborated by genetic data. However, in other cases, species are known only from few specimens and distinguished from congeners based on minute morphological differences alone. One such example is the genus *Pogonophryne* (Artedidraconidae), for which close to twenty species have been described within the last four decades (Eastman & Eakin, 2021), primarily on the basis of variation in the morphology of the mental barbel (*e.g*. Spodareva & Balushkin, 2014). This species richness within *Pogonophryne* could not be confirmed in a recent genetic analysis, which instead led to the synonymization of 24 out of 29 valid species (Parker, Dornburg, Struthers, Jones, & Near, 2022).

Outside *Pogonophryne*, the validity of species described in the genus *Channichthys* (Channichthyidae) remains particularly questionable. As many as nine species have been recognized, including the unicorn icefish (*C. rhinoceratus* Richardson, 1844), the red icefish (*C. rugosus* Regan, 1913), the sailfin pike (*C. velifer* Meisner, 1974), and six species described by Shandikov between 1995 and 2011 (Aelita icefish, *C. aelitae*; big-eyed icefish, *C. bospori*; pygmy icefish, *C. irinae*; charcoal icefish, *C. panticapaei*; green icefish, *C. mithridatis*; and robust icefish, *C. richardsoni*) (Shandikov, 1995a, 1995b, 2008, 2011) (Table 1). All species of the genus are endemic to the Kerguelen-Heard plateau and appear to share largely overlapping distributions (Shandikov, 2011), implying that they either represent a small radiation on their own, or that at least some of the described taxa are *de facto* morphs of one and the same species. The members of the genus are morphologically similar, with total lengths around 30–50 cm, a wide and spatulated snout, a tall first dorsal fin, and the rostral spine that gave its name to the the first described species, *C. rhinoceratus* (Richardson, 1844).

**Table 1.**
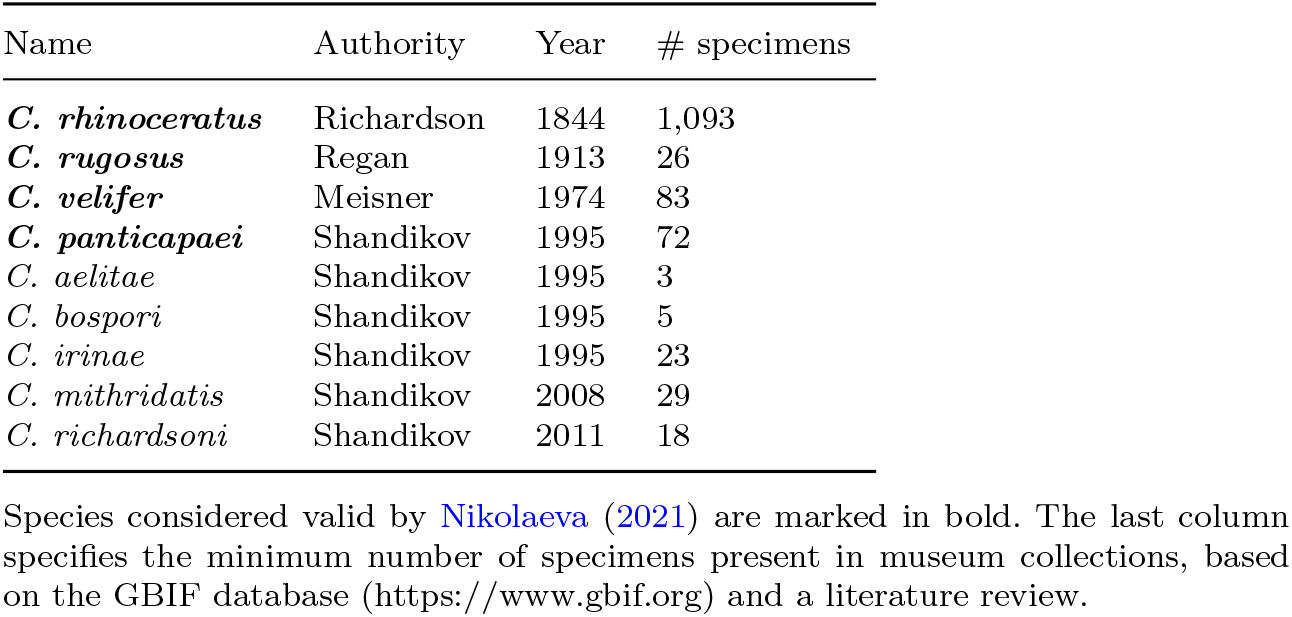
Species described in genus *Channichthys*

The second-oldest species, *C. rugosus*, was described on the basis of two specimens that were found to differ from the known *C. rhinoceratus* specimens in eye diameter, roughness of the head, the position of supraorbital edges, and the length of the maxillary (Regan, 1913). As additional specimens became available, the diagnosis of the two species became refined (Norman, 1937), but the lack of clearly species-defining traits led some authors to question their separation even before further members of the genus were described (Hureau, 1964). The third species to be described was *C. velifer* (Meisner, 1974), which was claimed to differ from *C. rhinoceratus* in the number of spines of the first dorsal fin and the presence of a single median series of bony plates on the posterior part of the body (Meisner, 1974). However, most of the specimens assigned to this new taxon were females, suggesting that sexual dimorphism could have explained the observed differences (Gon & Heemstra, 1990). The six remaining described species were added to the list to accommodate previously unseen combinations of ray numbers, interorbital width, fin membrane height, and a duplicated row of gill rakers, among others (Shandikov, 1995a, 1995b, 2008, 2011). In most cases, these species were described on the basis of few specimens only.

To address a felt demand for a “complete overhaul” (Duhamel, Gasco, & Davaine, 2005; Eastman & Eakin, 2021) of the systematics of the genus *Channichthys*, Nikolaeva and Balushkin began a series of investigations based on comprehensive comparisons of specimens in the collections of the Zoological Institute of the Russian Academy of Sciences in Saint Petersburg, the Ukranian National Museum of Natural History in Kyiv, and the British Natural History Museum in London. Their analyses indicated that the duplicate gill rakers observed in *C. rhinoceratus, C. panticapaei, C. bospori*, and *C. irinae*, but also in more distantly related icefishes, are a rather labile character (Balushkin & Nikolaeva, 2015), leading them to suggest the synonymization of the latter three *Channichthys* species (Nikolaeva, 2019). Their work further resulted in redescriptions of *C. velifer* (Nikolaeva & Balushkin, 2019), *C. rhinoceratus* (Nikolaeva, 2020), and *C. rugosus* (Nikolaeva, 2021), as well as the suggested synonymization of *C. aelitae, C. mithridatis*, and *C. richardsoni* with *C. rhinoceratus* (Nikolaeva, 2020).

Thus, according to Nikolaeva and Balushkin, the following four species are currently recognized in the genus *Channichthys*: *C. rhinoceratus, C. rugosus, C. velifer*, and *C. panticapaei*. The redescribed *C. rugosus* differs from *C. rhinoceratus* in four characters: greater height of the anterior dorsal fin, a fin membrane extending to the apexes of the longest rays, a narrower and concave interorbital space, and a more uniformly reddish body color (Nikolaeva, 2021). *Channichthys rugosus* can be further distinguished from *C. velifer* by numbers of fin rays in the first dorsal and the pectoral fin, bone plaques on the lateral line, and its coloration. Additionally, *C. panticapaei* was said to differ from *C. rugosus* in having a duplicated row of gill rakers and a more brownish-black coloration (Nikolaeva, 2021).

To complement the morphological analyses of *Channichthys* species and further test their validity, genetic data would be essential. Unfortunately, however, molecular information is so far only available for a single species, *C. rhinoceratus*. The genetic data available for this species include a set of ten nuclear markers commonly used in phylogenetic studies, cytochrome c oxidase I (*CO1*) barcodes (Smith et al., 2012), and the recently published complete sequence of a mitochondrial genome with a length of 17,408 bp (Andriyono et al., 2019), as well as restriction-site associated DNA (RAD) markers (Near et al., 2018). These sequences are available from the National Center for Biotechnology Information (NCBI). For the other three potentially valid species of the genus, the unavailability of sequence information is at least in part due to the rarity of suitable tissue samples. To the best of our knowledge no more than 3––72 specimens are present for these species in museum collections (Table 1). Moreover, most specimens of these species were caught decades ago and fixed in formalin, which leads to degradation and chemical modification of DNA, making the recovery of genetic information challenging. To overcome this limitation, protocols for DNA extraction and sequencing library preparation tailored for formalin-fixed specimens have recently become available and proved to be remarkably successful (Gansauge, Aximu-Petri, Nagel, & Meyer, 2020; Gansauge et al., 2017; Gould, Fritts-Penniman, & Gaisner, 2021).

Here, we apply recently developed methods to retrieve DNA sequences from a specimen of *C. rugosus* that was collected and formalin-fixed in 1974, and stored in 70 percent ethanol since. Modifications to a standard DNA extraction protocol from animal tissue maximise the yield of short DNA fragments, while application of a single-stranded DNA library preparation method (Gansauge et al., 2020, 2017) allows to convert even minute amounts of degraded DNA into sequencing libraries. We reconstruct the complete sequence of the specimen’s mitochondrial genome based on Illumina read data, and compare this sequence to the mitochondrial genome of *C. rhinoceratus* to assess the genetic divergence between these two species. We find that the two mitochondrial genomes are nearly identical, with only 27 nucleotide differences between them. Such close similarity lends support to the synonymity of the two species *C. rugosus* and suggests that, like *Pogonophryne*, *Channichthys* comprises fewer differentiated species than previously thought.

## 2 Materials and methods

### 2.1 Sampling

The *C. rugosus* specimen used for sequencing was collected on 28.06.1974 during voyage 7 of the Soviet scientific trawler ‘Skif’ (‘Cκиф’). It was obtained by bottom-trawling (trawl 188) on the Kerguelen shelf, at a depth of 115–120 m to the North-East of the Kerguelen Islands (48°34’1 S, 70°37’1 E). The specimen (ZIN 56294) has a standard length of 252 mm and is located in the collection of the Zoological Institute of the Russian Academy of Sciences in Saint Petersburg, Russia. It was formalin-fixed upon arrival in Saint Petersburg and remained in 40% formalin for several years before being transferred to 70% ethanol. The specimen had been identified as *C. rugosus* based on the height of its first dorsal fin, the shape of its interorbital space, its body coloration, numbers of fin rays, and the absence of a second row of gill rakers on the first gill arch (Nikolaeva, 2021).

### 2.2 DNA extraction and sequencing

A small piece of muscle tissue (5.8 mg dry weight) was dried in a vacuum centrifuge and immersed in lysis buffer (260 μL ATL buffer (Qiagen) and 40 μL Proteinase K [20 mg/mL]). Two extraction negative control reactions received the same lysis buffer, but did not contain sample. After 24 h incubation at 56°C, the lysates were centrifuged at 17,000 x g for 5 minutes and 300 μL supernatant mixed with 3000 μL Buffer PB (Qiagen). The mixtures were loaded onto Minelute silica columns (Qiagen) in steps of 600 μL, then washed twice with 600 μL Buffer PE (Qiagen). Centrifugation was carried out for 1 minute each at 8,000 x g, which is lower than recommended by the manufacturer, in order to increase the retention of short DNA fragments. The columns were dry spun 1 minute at 16’000 x g to remove residual wash buffer. To elute the DNA, 50 μL Buffer AE (Qiagen) were placed directly onto the silica membrane, incubated for 10 minutes and centrifuged at 16,000 x g for 2 minutes. DNA concentration was determined using 5 μL of the extract in a Qubit dsDNA High Sensitivity assay. The sample yielded 6.6 ng/μL, while the extraction negatives contained 0.0244 ng/μL or were below the detection threshold of 0.02 ng/μL, respectively. Eight μL of the extract (=52.8 ng DNA) and 30 μL each of the negative controls were then used to build Illumina sequencing libraries by single-stranded DNA library preparation as described by Gansauge et al. (2020, 2017), using the same reagents as listed in Gansauge et al. (2020). Briefly, the DNA extract and two extraction negative controls, along with one library negative control (water) and one library positive control (0.1 pmol of oligonucleotide CL104) were used for five separate library build reactions. Furthermore, all reactions received 10 amol CL104 as internal control. Samples were dephosphorylated and the 3’-adapter (TL181/TL110) attached by splinted ligation. Adapter molecules were bound to streptavidin-coated magnetic beads which were carried through the reactions and washed after each enzymatic reaction. An adapter-complementary primer (CL128) was used to prime the fill-in reaction, creating 5’-blunt-ended double stranded molecules. Another ligation attached the second adapter (CL53/TL178). Libraries were eluted into 50 μL Tween-20 supplemented Tris-EDTA buffer (i.e. TET buffer) by heat denaturation. 1 μL each of a 1:50 dilution of libraries was used in two qPCR assays to determine control molecule numbers and required PCR-cycle numbers for amplification. The remaining 49 μL of libraries, with the exception of the positive library control, were then uniquely dual indexed (7 bp index length) (Kircher, Sawyer, & Meyer, 2012) and amplified until the end of the exponential phase, then purified using the Minelute PCR purification kit (Qiagen). Libraries were pooled and size-selected to 160–250 bp on a Blue Pippin instrument using a 3% cassette with internal markers. The selected size fraction was measured with both, the Qubit dsDNA High Sensitivity assay and the Tapestation HS1000 assay to adjust the input molarity for the sequencing run. The pool was then sequenced on an Illumina NextSeq500 instrument using a high-output single-end 75 bp read length kit with custom primers for read 1 (CL72) and index 2 (Gesaffelstein) (Paijmans et al., 2017).

Raw read files were demultiplexed using bcl2fastq v.2.19.1 (https://support.illumina.com/sequencing/sequencing_software/bcl2fastq-conversion-software.html), allowing for a maximum combined distance of 1 between barcodes, and saved to fastq format. In the same step, prevalent adapter sequences were trimmed from the ends of reads, and reads under 20 bp of length were omitted.

To ensure the presence of endogenous reads, we matched each read to the NCBI non-redundant (NR) sequence database (downloaded on 12 February 2022) with the BLASTX algorithm as implemented in Diamond v.2.0.4 (Altschul, Gish, Miller, Myers, & Lipman, 1990; Buchfink, Xie, & Huson, 2015). The resulting taxon assignments were plotted with MEGAN v.6.21.4 (Huson et al., 2016).

### 2.3 Reference-based sequence analysis

We mapped reads to the *C. rhinoceratus* mitochondrial genome (Genbank accession number NC_057120) using BWA v.0.7.17 (Li & Durbin, 2010), with its “aln” algorithm and the maximum fraction of missing alignments set to 0.05. The *C. rhinoceratus* mitochondrial genome was obtained from an individual that was collected in 2018 from Antarctic Subarea 58.5.2 (Heard Island and McDonald Island; Sapto Andriyono and Hyun-Woo Kim, priv. comm.). Resulting alignments were filtered to a Phred-scaled mapping quality of 25 or higher with Samtools v.1.9 (Li et al., 2009). Duplicated reads, e.g. from PCR duplicates, were flagged with Picard’s v.2.21.3 (http://broadinstitute.github.io/picard/) MarkDuplicate function and filtered with Samtools. Sites with a minimum read depth of 3 were consensus called using ANGSD v.0.933 (Korneliussen, Albrechtsen, & Nielsen, 2014). To recover the terminal positions of the sequence, the first 200 bp of the reference were cut and appended, and the mapping repeated. Regions of low-complexity and larger repeats exceeding the read length were edited manually or replaced with “N” for the length of the reference, as described in the Results section. The degradation state of DNA was assessed by inspection of the distribution of read lengths and analysis of substitution frequencies per base position within reads, using mapDamage v.2.0.8 (Jónsson, Ginolhac, Schubert, Johnson, & Orlando, 2013).

### 2.4 Reference-independent sequence analysis

To exclude potential reference bias, we also performed analyses of the mitochondrial genome of *C. rugosus* based on *de novo* assembly. We performed local assembly of individual mitochondrial markers with aTRAM v.2.0 (Allen, LaFrance, Folk, Johnson, & Guralnick, 2018). As queries, we used nucleotide and protein sequences from all mitochondrial genomes available on NCBI. To identify these notothenioid mitochondrial genomes on NCBI, we used the search string ““Notothenioidei”[Organism] AND (“mitochondrial”[Title] OR “mitochondrion”[Title]) AND “complete genome”[Title]” on 4 December 2021. This set of mitochondrial genomes included the one for *Channichthys rhinoceratus* and 40 other unique mitochondrial genomes. From each of these 41 mitochondrial genomes, we extracted each gene (rRNA, tRNA, or protein-coding) in nucleotide format, and protein-coding features in amino-acid format, and used all of these in aTRAM analyses. All aTRAM analyses were performed separately with seven different e-value thresholds (1e-2, 1e-3, 1e-4, 1e-5, 1e-6, 1e-8, 1e-10) for the BLASTN and TBLASTN v.2.10.1 (Altschul et al., 1990) searches that aTRAM runs internally. As the assembler tool internally employed by aTRAM, we selected Trinity v.2.10.0 (Grabherr et al., 2011). Each aTRAM analysis was continued for 20 iterations.

All contigs produced by aTRAM were jointly used as input for a second assembler tool, MIRA v.4.9.6 (Chevreux, Wetter, & Suhai, 1999). We set MIRA’s “nasty repeat ratio” to 25 (“-KS:nrr=25”), the “maximum megahub ratio” to 40 (“-SK:mmhr=40”), specified the lack of quality information (“-AS:epoq=no”), and turned off the checks for average coverage (“-NW:cac=no”) and maximum read name length (“-NW:cmrnl=warn”), according to the format of the input data. The contigs produced by MIRA (or their reverse complements) were then individually aligned to the mitochondrial genome for *C. rhinoceratus* with MAFFT v.7.470, using a gap opening penalty of 2, a gap extension penalty of 1, and the program’s “6merpair” and “addfragments” options.

### 2.5 Comparative analyses

The reference-based and reference-independent mitochondrial genome sequences for *C. rugosus* were compared visually using AliView v.1.2.6 (Larsson, 2014). Nucleotide sequences of the 13 protein-coding genes were extracted from the mitochondrial genome of *C. rugosus* and all other notothenioid mitochondrial genomes and aligned per gene with MAFFT. The 13 alignments were concatenated and a distance matrix was calculated from the concatenated alignment with the Python script “convert.py” (available from GitHub: https://github.com/mmatschiner/supergenes/blob/main/gadidae_phylogenomics/src/convert.py), ignoring all sites with missing data. The concatenated alignment was further used for maximum-likelihood phylogenetic inference with IQ-TREE v.2.1.2 (Minh et al., 2020), applying the program’s automated substitution model selection and 1000 ultrafast bootstrap (Minh, Nguyen, & von Haeseler, 2013) iterations.

To gain a more complete view of sequence variation in *Channichthys*, we took advantage of the *CO1* barcode sequences available on NCBI (Smith et al., 2012). We downloaded these sequences and aligned them together with the homologous sequences extracted from the mitochondrial genomes of *C. rhinoceratus* and *C. rugosus*, using MAFFT. To illustrate the *CO1* sequence variation in *Channichthys*, we first applied maximum-likelihood phylogenetic inference with IQ-TREE and then used the estimated phylogeny jointly with the sequence alignment to draw a haplotype genealogy graph with the program Fitchi v.1.1.4 (Matschiner, 2016).

## 3 Results

### 3.1 Sequencing

For the *C. rugosus* specimen, a total of 63,158,791 reads passed initial trimming and length filtering. For extraction negative controls 1 and 2 (exneg 1, exneg 2), and library negative control (libneg), the number of reads were 22,393,754, 21,343,774 and 3,080,057, respectively. Analysis with Diamond and MEGAN assigned a substantial proportion of the reads to Notothenioidei and confirmed that these were endogenous (Fig. 4).

### 3.2 Reference-based sequence analysis

Mapping to the *C. rhinoceratus* mitochondrial genome generated 48,313 hits, corresponding to 0.0765% of total reads. After filtering duplicates, 18,687 unique reads were used for further analysis. These reads had an average length of 32.98 bp, with only 0.1% of reads being 70 bp or longer. Figure 2 shows the distribution of read lengths of hits. Negative controls produced 5 (exneg 1), 5 (exneg 2), and 1 unique hit (libneg), which weren’t analysed further. The resulting coverage had an average depth of 35.4 and extended over 98.1% of the reference. The alignment showed two apparent gaps in the D-loop and spurious alignments directly adjacent to them, therefore positions 15,192–15,294 and 16,856–17,262, respectively, were called as ”N” for the length of the reference. A 14-mer C repeat at reference positions 15,986–15,999 was spanned by two reads only, both indicating an indel with one fewer repeat in *C. rugosus*, which was called manually. Mapped reads showed an elevated rate of C to T changes in both their 3’- and 5’-ends, a pattern of cytosine deamination that is typical for degraded DNA (Fig. 3).

**Figure 1.**
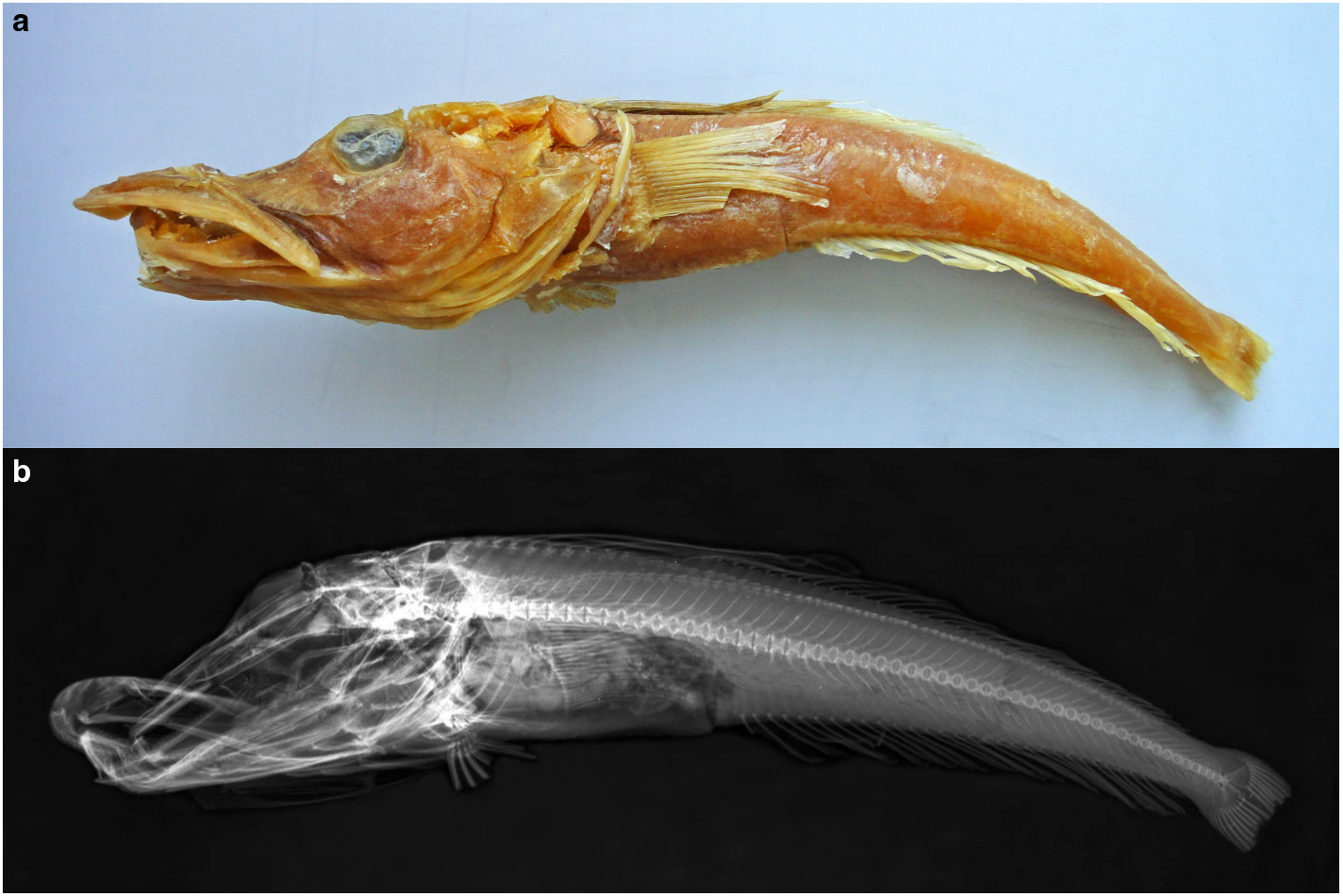
*Channichthys rugosus* specimen ZIN 56294. Photograph (a) and x-ray image (b) of the *C. rugosus* specimen used for DNA extraction. The specimen has a standard length of 252 mm.

**Figure 2.**
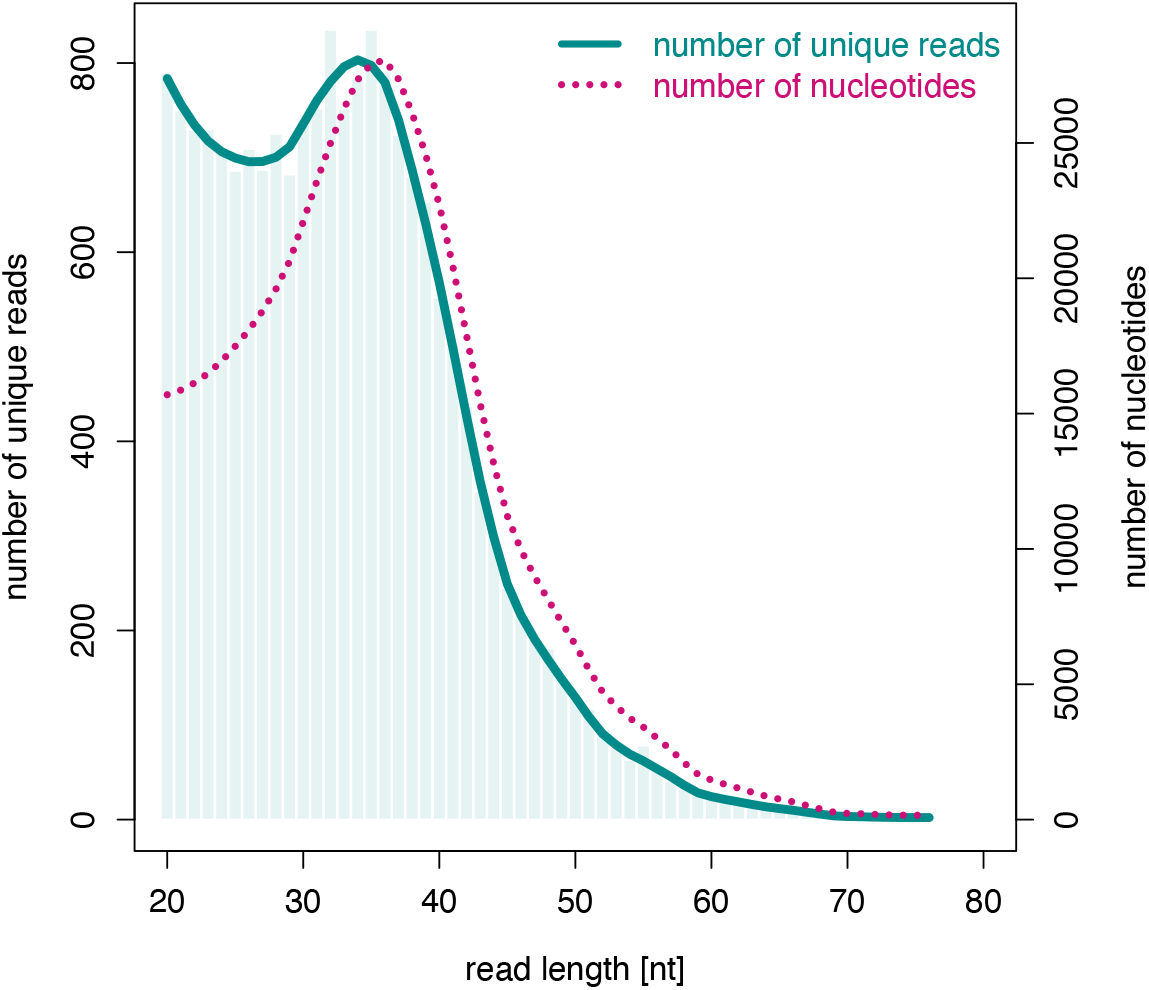
Distribution of read lengths of hits to the *C. rhinoceratus* mitochondrial genome. The solid line and bars indicate the number of hits for a given read length, the dotted line indicates the amount of data in nucleotides gathered from reads of a given length.

**Figure 3.**
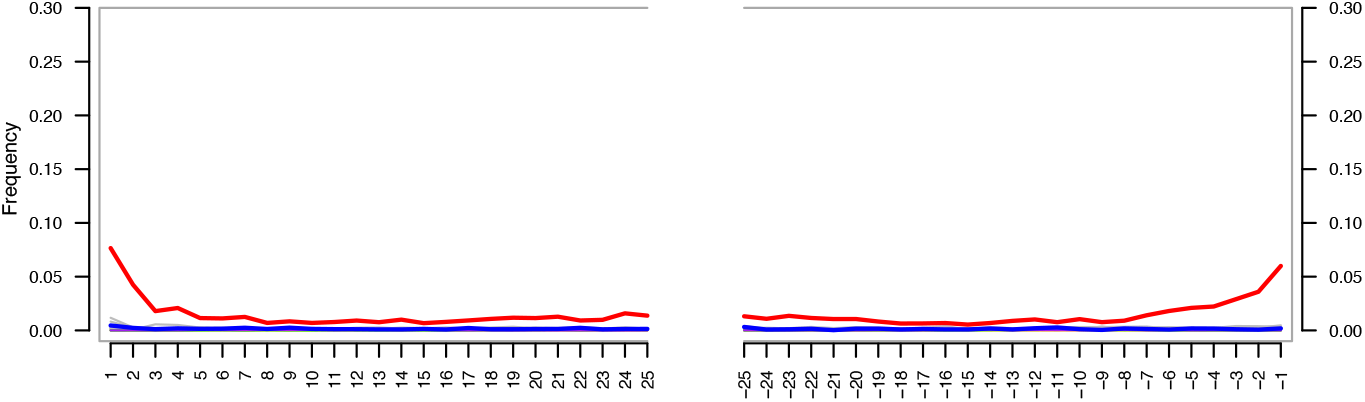
Frequency of post-mortem C to T changes by nucleotide position within reads. Deaminated cytosines are being read as thymine and are especially prevalent at molecule ends. Red line: relative frequency of T by position in read from 5’-end (left) and 3’-end (right), blue line: relative frequency of A. Here, the deamination signal manifests only as C to T, and not G to A, because single-stranded, rather than double-stranded DNA library preparation was used.

**Figure 4.**
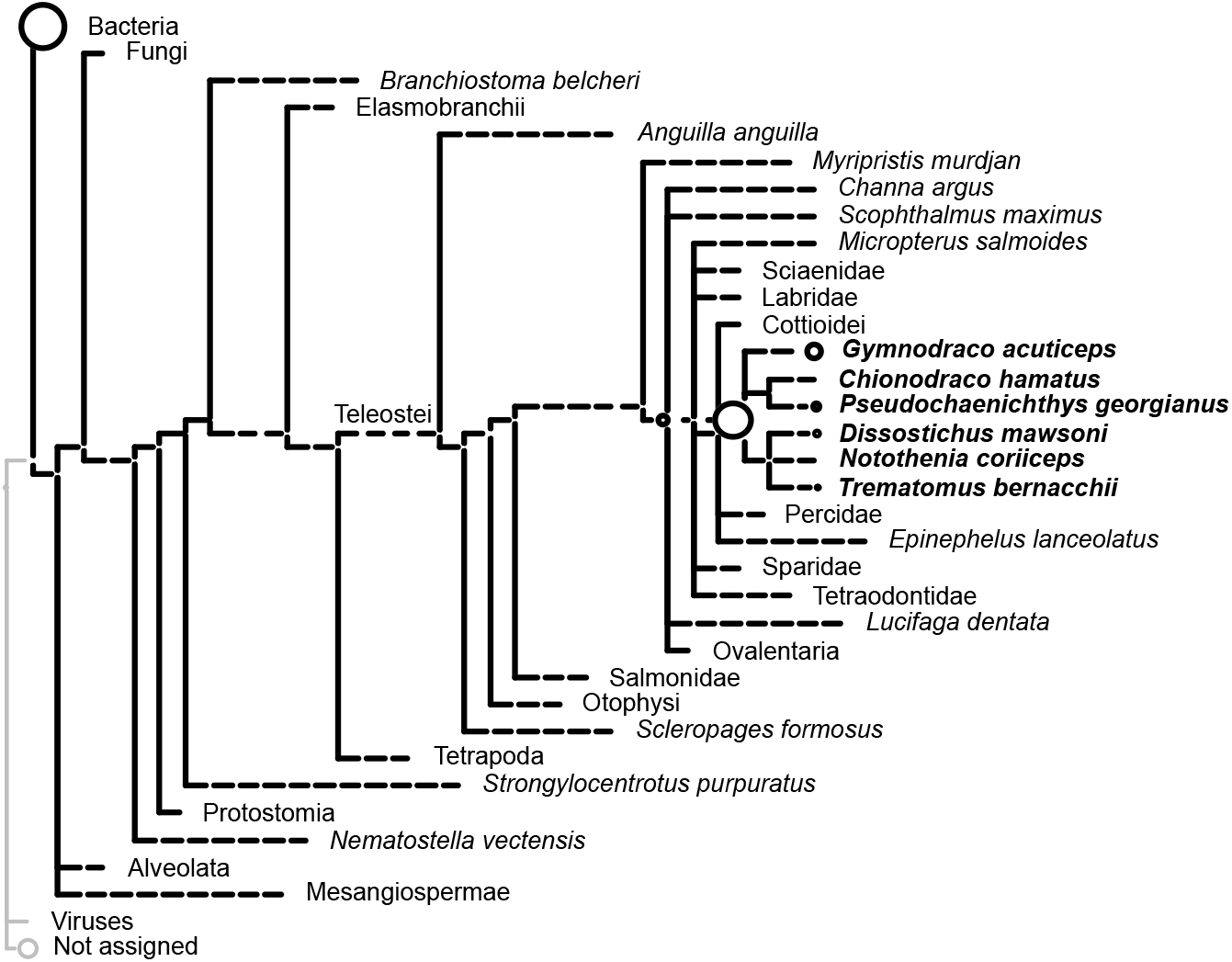
Taxonomic assignment of individual reads. Reads were mapped to the NCBI non-redundant (NR) database. The sizes of circles on internal and terminal nodes are proportional to the numbers of reads mapping to the corresponding taxon; a single read mapped to taxa without visible circles.

### 3.3 Reference-independent sequence analysis

Assembly with MIRA produced 14 contigs with lengths between 317 and 1471 basepairs (bp) (mean length: 677.4 bp). One of these contigs overlapped fully with another one, and two other contigs overlapped by 13 bp. The total length of the mitochondrial genome covered by these contigs was 9,849 bp. No nucleotide differences were observed between overlapping contigs.

### 3.4 Comparative analyses

Comparison of the reference-based and the reference-independent mitochondrial genome sequences showed that these were completely identical across the 9,849 bp covered by both. Compared to the mitochondrial genome of *C. rhinoceratus*, the reference-based sequence differed at 27 sites (Table 2). Seventeen of these nucleotide differences were within regions also covered by the reference-independent sequence for *C. rugosus*, and all of these were confirmed by that sequence. The differences between the *C. rhinoceratus* and *C. rugosus* mitochondrial genomes included five transitions (two A/C, one G/C, one G/T, and one T/G substitution) and 21 transversions (seven A/G, two C/T, six G/A, and six T/C substitutions), as well as one C/- indel. The nucleotide differences were distributed unevenly across the mitochondrial genome and mostly found in the *ND6* gene and the D-loop region (Fig. 5). Compared to the mitochondrial genome-wide background of 0.0011 substitutions per bp, the *ND6*/D-loop region had an elevated divergence of 0.0065 substitutions per bp.

**Table 2.** Nucleotide substitutions and indel between *Channichthys* mitochondrial genomes

| Site | Nucleotide substitution | Codon position | Codon substitution | Amino-acid substitution |
| --- | --- | --- | --- | --- |
| 1379 | a/g |  |  |  |
| 1552 | c/t |  |  |  |
| 2784 | a/g |  |  |  |
| 2893 | a/g | 3 | cta/ctg |  |
| 3145 | g/c | 3 | ctg/ctc |  |
| 4165 | t/c | 3 | att/atc |  |
| 4410 | g/a | 2 | ggc/gac | G/D |
| 4994 | a/g | 1 | acc/gcc | T/A |
| 5857 | g/a | 3 | tgg/tga |  |
| 7586 | a/g | 3 | gaa/gag |  |
| 8650 | g/a | 2 | ggc/gac | G/A |
| 9234 | t/c | 3 | ctt/ctc |  |
| 10761 | g/a | 3 | acg/aca |  |
| 11617 | t/c | 1 | ttt/ctt | F/L |
| 13006 | t/c | 3 | gct/gcc |  |
| 13832 | g/a | 1 | gag/aag | E/L |
| 14550 | a/c | 3 | gga/ggc |  |
| 15568 | t/g | 2 | gag/gcg | E/A |
| 15647 | a/c | 1 | tca/gca | S/A |
| 15831 | a/g | 3 | gct/gcc |  |
| 15882 | a/g | 3 | tgt/tgc |  |
| 15999 | c/- |  |  |  |
| 16049 | c/t |  |  |  |
| 16138 | g/t |  |  |  |
| 16147 | g/a |  |  |  |
| 16477 | t/c |  |  |  |
| 17347 | t/c |  |  |  |

**Figure 5.**
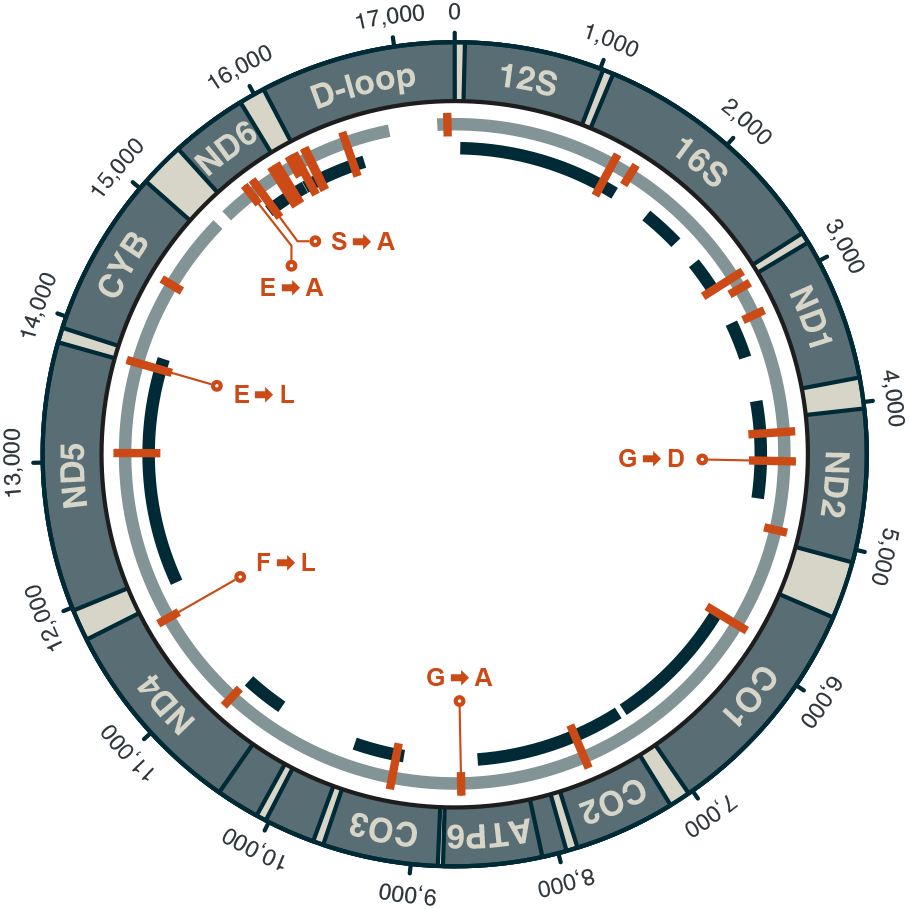
Mitochondrial genome of *C. rugosus*. Coordinates on the outside of the circle are given in units of basepairs. The gray inner circle indicates the nearly full coverage achieved in the reference-based analysis. Two gaps with a total length of 510 bp remain in this sequence between positions 15,192 and 15,294, and between positions 16,856 and 17,262. The black fragments inside the gray circle show the positions of contigs from the reference-independent approach. Substitutions compared to the mitochondrial genome of *C. rhinoceratus* (NCBI accession NC_057120) are marked in orange, and non-synonymous substitutions are labelled with the resulting amino-acid change.

Phylogenetic inference with IQ-TREE based on protein-coding mitochondrial sequences grouped the two *Channichthys* genomes with full bootstrap support. The two taxa were separated only by very short branches that had a combined length of 0.0012 substitutions per bp. The *Channichthys* mitochondrial genomes were most similar to those of *Chionobathyscus dewitti* and *Cryodraco antarcticus*, with which they formed a clade that also received full bootstrap support. This clade appeared as the sister group to a clade formed by the mitochondrial genomes of *Chaenodraco wilsoni* and three members of the genus *Chionodraco*, albeit with low bootstrap support of 70% (Fig. 6).

**Figure 6.**
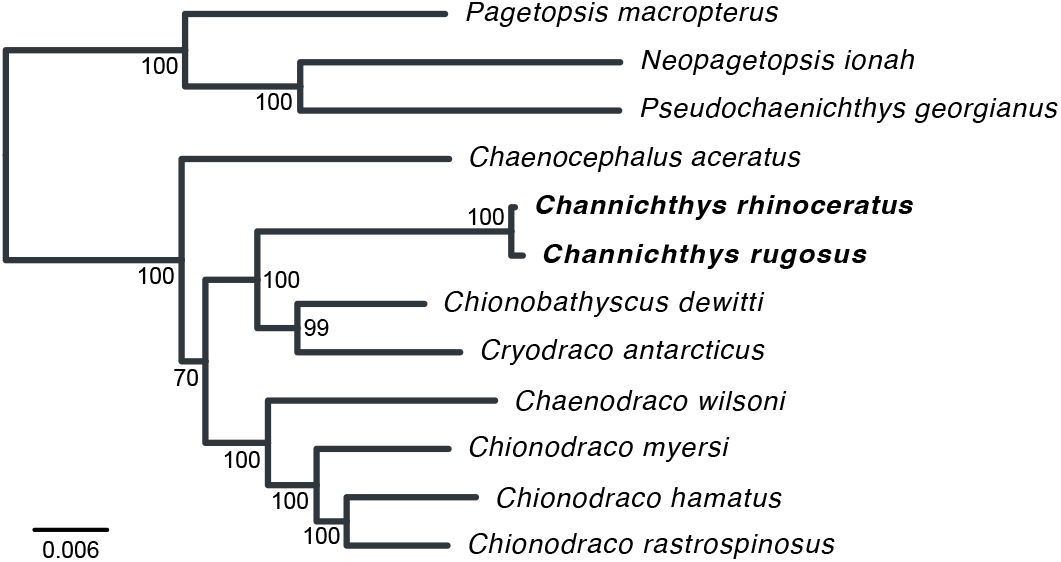
Mitochondrial phylogeny of Channichthyidae. Maximum-likelihood phylogeny of all available mitochondrial genomes for Channichthyidae. Short branches connect *Channichthys rhinoceratus* and *C. rugosus*.

The haplotype genealogy graph for 14 available *Channichthys CO1* sequences showed that eleven of them shared the same haplotype. Three other haplotypes differed by a single substitution from the majority haplotype and were each represented by a single individual. One of these three private haplotypes was found found in the published mitochondrial genome sequence for *C. rhinoceratus*, while the *C. rugosus* mitochondrial genome had the majority haplotype (Fig. 7).

**Figure 7.**
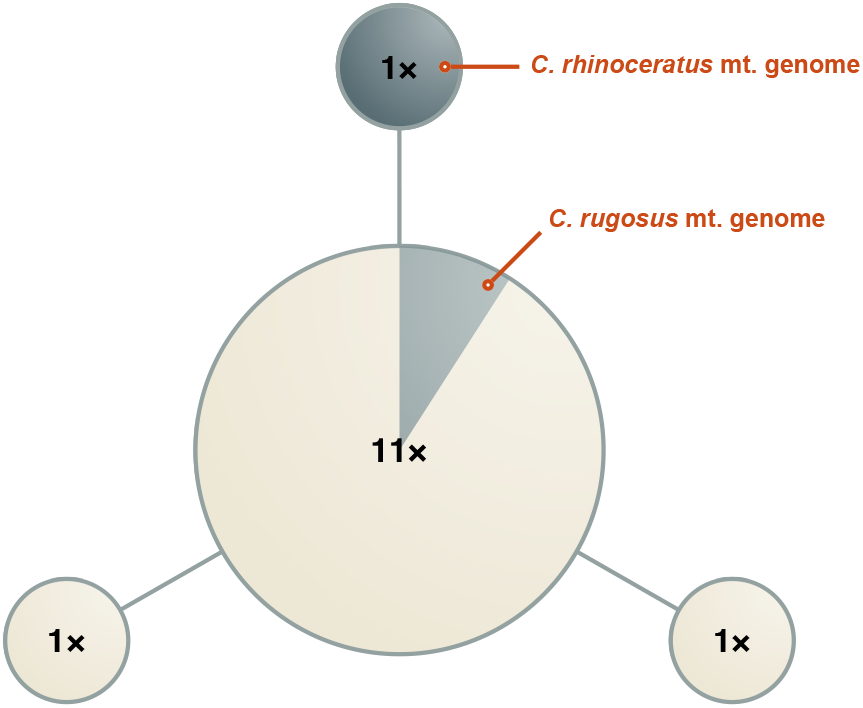
Haplotype genealogy graph for *CO1*. Circles represent four distinct *Channichthys CO1* haplotypes, found in 13 *C. rhinoceratus* and one *C. rugosus* individuals. Radii are drawn according to the number of individuals with that haplotype (indicated with labels on circles). Edges represent substitutions. Each of the three edges has a length of one substitution. mt., mitochondrial.

## 4 Discussion

Our comparison of the mitochondrial genomes of *C. rhinoceratus* and *C. rugosus* showed that these two mitochondrial genomes are highly similar, with only one indel, 26 nucleotide substitutions and six amino-acid substitutions between them. The sequence divergence is 0.16%, and most of this divergence is concentrated in the *ND6*/D-loop region where the divergence reaches 0.65%. Despite very short read lengths, reconstruction of *C. rugosus*’ mitochondrial genome was possible, except for two loci in the ND6/D-loop region. Those are most likely repeats that were too large to be spanned by single reads and could not be mapped correctly. While larger rearrangements of nothotenioid mitochondrial genomes have been reported by Papetti et al. (2021), gene order was shown to be identical for eight of the species in figure 6 (all except *Chionobathyscus dewitti, Cryodraco antarcticus*, and *Channichthys* sp.). Hence, while possible, the gaps in the alignments are unlikely to be caused by rearrangements or larger indels, and are probably artifacts due to short read length. This highlights a shortcoming of the use of degraded DNA, where read lengths are usually limited by DNA molecule lengths rather than by sequencing technology, making the detection of structural variants challenging.

Analysis of post-mortem damage shows a robust pattern of cytosine deamination in single-stranded overhangs at molecule ends, manifesting as C to T changes in the sequencing data. Given the appreciable depth of coverage, however, we consider it very unlikely that any position was called erroneously due to this damage. Here, only C to T changes are seen as we used single-stranded DNA library preparation. In double-stranded DNA library preparation, deaminated cytosines would also be apparent as G to A changes in the sequencing data. The presence of this deamination signal can be interpreted as evidence for the authenticity of the sequences, as modern contamination would not show it. However, this is more relevant for ancient samples, for which the age of endogenous DNA and *ex situ* contamination would be very different, and for cases where contamination is more likely to be confused with endogenous sequences, such as ancient human samples. Here, contamination is unlikely to affect our results, as negative controls didn’t produce concerning numbers of reads mapping to the reference, and no other samples or DNA were handled in the laboratory environment which couldn’t be readily distinguished from our sample.

The available molecular data for the genus *Channichthys* does not allow us to perform a formal genetic species delimitation analysis as was recently done for *Pogonophryne* (Parker et al., 2022). Nevertheless, a comparison of the divergence between the two *Channichthys* mitochondrial genomes with the levels of between- and within-species sequence divergence in other notothenioid fishes can inform about the existence of one or several species within the genus. As shown in Fig. 6, the divergence within *Channichthys* is far smaller than that between any other pairs of *Channichthyidae* species. Besides *Channichthys rhinoceratus* and *C. rugosus*, the most closely-related species (judged on the basis of their mitochondrial genomes) appear to be *Chionodraco hamatus* and *Chionodraco rastrospinosus*. However, despite their recent divergence around a million years ago (Colombo, Damerau, Hanel, Salzburger, & Matschiner, 2015), the mitochondrial genomes of these two species are connected by a total branch length of 0.019 substitutions per bp, around 16 times as long as the branches connecting the two *Channichthys* mitochondrial genomes. Of note, the two *Chionodraco* species appear not to be fully separated, given that hybrids and signals of introgression have been observed (Schiavon et al., 2021).

On the other hand, the divergence between the two *Channichthys* mitochondrial genomes is comparable to the within-species divergence in other notothenioid species. Mitochondrial genomes of two individuals are available for five notothenioid species: *Trematomus borchgrevinki*, *Notothenia coriiceps*, *Notothenia rossi*, *Chaenodraco wilsoni*, and *Chionodraco hamatus*. These two genomes per species differ by 16–92 substitutions and 3–22 indels across the mitochondrial genome, or by 11–69 substitutions and 2-10 indels when the *ND6* / D-loop region is excluded (Table 3). The divergence of the two *Channichthys* mitochondrial genomes is close to or below the lower ends of these ranges, considering that we found these two genomes to differ in 26 nucleotide substitutions (17 outside the *ND6* / D-loop region) and no indels were observed. In contrast, the two most-closely related pairs of sister species within Channichthyidae (*Chionodraco hamatus*, *Chionodraco rastrospinosus*, and *Chionobathyscus dewitti, Cryodraco antarcticus*; Fig. 2) differ by 421–540 substitutions and 25–34 indels (253–272 substitutions and 6–11 indels when the *ND6* / D-loop region is excluded; Table 3). Thus, the divergence of the mitochondrial genomes of *Channichthys rhinoceratus* and *C. rugosus* is consistent with the existence of a single species within the genus *Channichthys* (Eastman & Eakin, 2021). If this result should be confirmed by further molecular data, *Channichthys rugosus* may need to be synonymized with *C. rhinoceratus*.

**Table 3.**
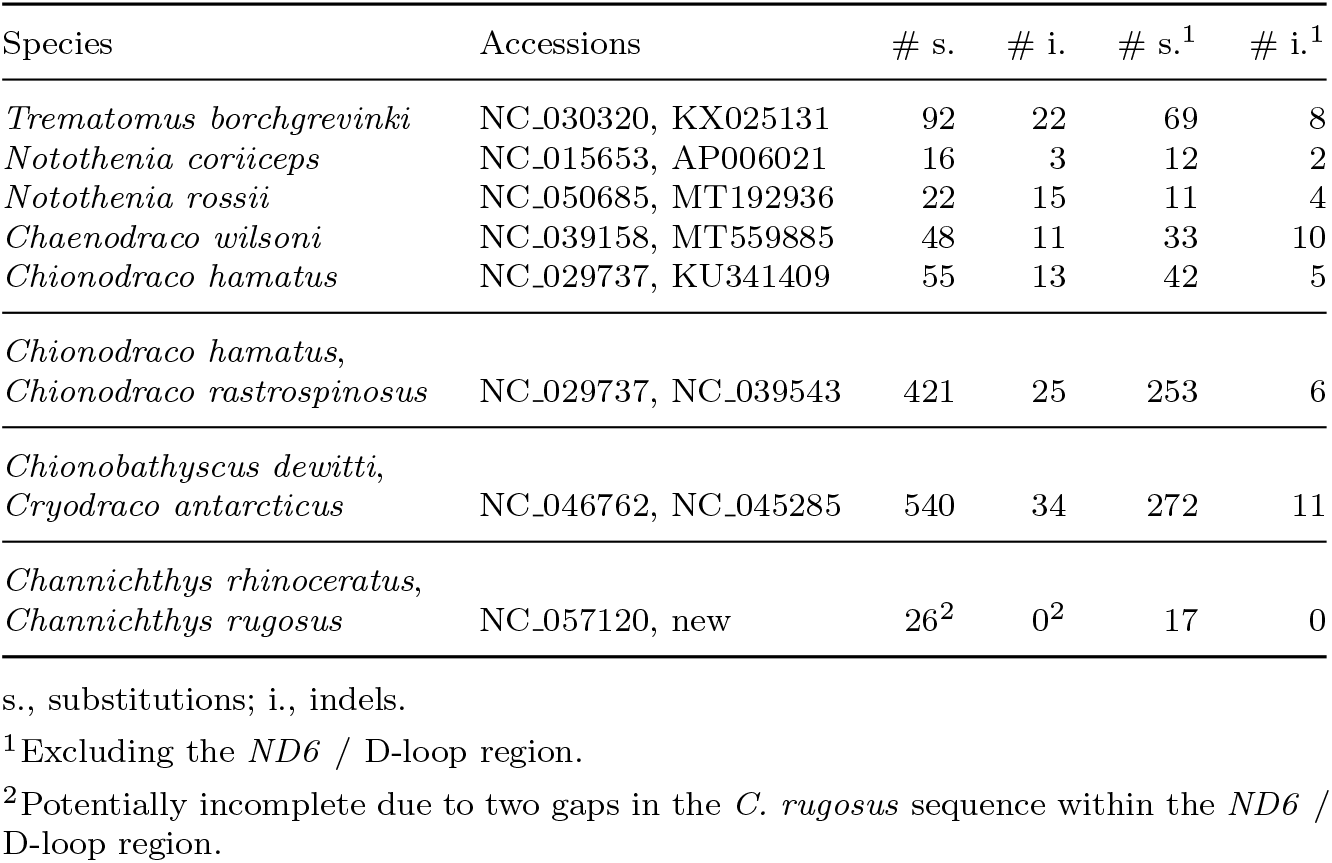
Within-species mitochondrial sequence divergence in Notothenioidei and between-species divergence for closely-related Channichthyidae

Low divergence in mitochondrial genome sequence can be indicative of a close – e.g. intraspecific – phylogenetic relationship, but could also be due to alternative scenarios. Mitochondrial capture, where one lineage fixes the mitochondrial genome received from another by introgression, can result in two valid, reproductively largely isolated species with appreciable nuclear genome divergence having no or little mitochondrial genome divergence. This phenomenon can also apply to large, rapidly diversifying clades where many species have arisen from nuclear genomic diversity created through hybridisation of divergent ancestral lineages, for example in Lake Victoria’s haplochromine cichlids (Meier et al., 2017). In that case low mitochondrial diversity and rampant haplotype sharing would conceal a large species diversity. It would need to be tested using genome-wide nuclear data if such a scenario applies to *Channichthys*, rather than a previous overestimation of species diversity in the genus.

Genetic data can be vital to corroborate or reject taxonomic assessments based on morphology and to more accurately estimate organismal diversity. The fixation of specimens with formalin, which severely hampers genetic analyses, has long been recognized, and indeed lamented, as a major obstacle for tapping the theoretically vast potential of museum collections for addressing long standing questions in systematics, taxonomy, evolutionary and conservation biology (Card, Shapiro, Giribet, Moritz, & Edwards, 2021; Raxworthy & Smith, 2021). Time-structured samples, including extinct species, can reveal details about the recent decline of biodiversity. Specimens from places that are difficult to access could fill gaps in studies otherwise relying on a smaller geographic sampling. Given this potential it is unsurprising that the research community continues to undertake great efforts in refining methodology to maximise the data yield from wet–collection specimens (Campos & Gilbert, 2012; Hahn et al., 2021; Hykin, Bi, & Mcguire, 2015; Schander & Kenneth, 2003; Straube et al., 2021). Here, we demonstrated that recently proposed methods that build on advances in the study of ancient DNA and high-throughput shotgun sequencing can recover usable genetic data from a formalin-fixed fish specimen. The short molecule lengths and very low DNA amounts recovered from such samples mandate the use of specialised, sensitive methods, such as lower centrifugation speeds and greater excess of binding buffer in DNA extraction (Dabney et al., 2013). The chemical cross-linking of DNA with DNA and proteins requires a harsher treatment of the tissue sample during lysis, for example by using high amounts of proteinase, as done here. Subsequently, the use of very efficient single-stranded DNA library preparation can convert appreciable numbers of molecules even if chemical damage and cross-linking are prevalent. Caution should be taken when analysing such low-yield samples, as levels of contamination regarded minor in other circumstances can obscure signal from the target or lead to incorrect interpretations. Working in a clean laboratory that is dedicated to the analysis of low-biomass, degraded samples is therefore recommended, as well as including appropriate negative controls to monitor contamination. This extra effort, however, is greatly rewarded when open questions can be addressed with otherwise unobtainable data.

## Code Availability

Analysis code is available from GitHub (https://github.com/mmatschiner/unsequenced).

## Acknowledgments

We thank Arcady V. Balushkin for providing the *C. rugosus* specimen and Marcelo Sánchez-Villagra for financially supporting the sequencing of its mitochondrial genome. Sapto Andriyono and Hyun-Woo Kim provided helpful information about the *C. rhinoceratus* mitochondrial genome. Mark Lever (ETH Zurich) kindly provided lab space. The Genetic Diversity Centre (ETH Zurich) provided access to laboratory and HPC facilities. The Functional Genomics Centre Zürich (ETH and University of Zurich) provided assistance with sequencing.

## Author Contributions

M.Mu. performed molecular lab work, bioinformatic analyses, and contributed to the manuscript. E.N. initiated the study and contributed the tissue sample of *C. rugosus*. L.R. established the collaboration. M.Ma. performed bioinformatic analyses and wrote most of the manuscript. All authors read and approved the final version of the manuscript.

## Funding

M. Muschick was supported by the SNSF Sinergia grant CRSII5_183566. M. Matschiner was supported by the Norwegian Research Council with FRIPRO grant 275869. E. Nikolaeva was supported by a research programme of the Zoological Institute of the Russian Academy of Sciences (project number 122031100285-3).

## Declarations

### Conflict of interest

The authors declare that no competing interest exists, as well as that there is no financial support or relationships that may pose any kind of conflict. Likewise, the authors declare that contributed to the text, agreed with its content and approved it for submission.

## Notes

### Competing Interest Statement

The authors have declared no competing interest.

